# Enzyme promiscuous profiles for protein sequence and reaction annotation

**DOI:** 10.1101/2023.09.13.557547

**Authors:** Homa MohammadiPeyhani, Anastasia Sveshnikova, Ljubisa Miskovic, Vassily Hatzimanikatis

**Author notes:** These authors contributed equally (Homa MohammadiPeyhani, Anastasia Sveshnikova).

## Abstract

Novel sequencing techniques and biochemical pathway prediction resources provide a wealth of data on novel proteins and computationally predicted enzymatic reactions. Accurate matching of protein sequences to enzymatic activities is crucial for advancing synthetic biology and metabolic engineering efforts. Here we present BridgIT+, a computational workflow that accounts for enzyme promiscuity and accurately predicts protein-reaction and reaction-protein associations. BridgIT+ builds upon the promiscuity-based method for annotating orphan and novel reactions with enzymatic activities, BridgIT, and utilizes position-specific scoring matrices (PSSM). The framework uses sequence alignment and enzyme promiscuity predictions to analyze protein sequences, identify sequence patterns, and create promiscuous protein sequence profiles for each reaction. These profiles allow us to predict the protein sequences most likely involved in the reaction. We showcase BridgIT+ by annotating (i) computationally predicted reactions with proteins and (ii) unannotated proteins of *E. coli* proteome with enzymatic functions. We demonstrated the performance of BridgIT+ on several biochemical assays and compared it to three current state-of-the-art methods for matching proteins and reactions. We anticipate that the proposed conceptual framework will enhance our understanding of gene-protein-reaction relations and advance biological sequence and reaction annotation in biology and synthetic biology studies.

## 1.2 Introduction

While the number of fully sequenced genomes rapidly increases, their functional annotation lags behind^1^. It has been estimated that 30% of unannotated sequences have a metabolic function, indicating the critical knowledge gap in our understanding of enzymes and their role in cellular metabolism^2^. This gap exists because functional annotation of uncharacterized enzymes requires extensive *in vitro* and *in vivo* experiments. Computational methods (Supplementary material 1) can significantly reduce the time and cost of this process.

For enzymatic function annotation, researchers use enzymatic function descriptors^3^, such as the Enzyme Commission (EC) number, which systematically classifies enzymes based on the associated biochemical reactions. Current computational methods focus on inferring the EC number from a sequence using mainly two approaches: (i) data-driven approaches based on machine learning and (ii) homology-based approaches based on the available biochemical knowledge^4^. In recent studies, machine learning (ML) has been successfully used in all stages of protein annotation, from structural prediction to functional analysis^5^. For example, DeepEC correctly annotated protein sequences with EC numbers with high precision and sensitivity^6^. However, ML-driven results often need more interpretability and heavily depend on the dataset structure and training parameters^7^. In comparison, homology-based methods use a rational approach to identify evolutionarily conserved sequence patterns often representing the protein function. For example, PRIAM employs EC-based enzymatic profiles to account for the relationship between chemistry and enzyme sequences and improve the functional annotation of uncharacterized sequences^4^. The basic premise of protein homology is that similar sequences are derived from a common ancestor and have the same function^8^. However, this assumption cannot explain the functional similarity of orthologs or the different functionality between paralogs^4^. Another major limitation of current homology-based methods is that they can only functionally annotate a protein with similarity to other sequenced proteins^3^. Therefore, we need new approaches to broaden the search space for functional annotation of proteins.^6^

The links between chemistry and protein sequences are intricate, and phenomena such as the substrate promiscuity of the enzymes^9^ should be considered in the enzyme annotation process. This idea was put forward in the BridgIT^10^ method, showing that enzyme promiscuity is essential for discovering the secondary functions of enzymes. BridgIT uses reactive-site-specific fingerprints, originating from the expert-curated biochemical reaction rules^11–14^, to match reactions based on structural and functional similarity and predict the EC class for orphan and novel reactions.

Here, we present BridgIT+, an approach that goes beyond BridgIT capabilities and directly links orphan protein sequences and orphan reactions. In contrast to prominent enzyme annotation tools DeepEC and PRIAM that assign the EC class to protein sequences, BridgIT+ assigns *reactions* to protein sequences. Indeed, it captures enzymatic functions based on the reaction mechanism similarity rather than relying on the EC classification. This way, BridgIT+ overcomes limitations imposed by the EC classification such as misclassification, unclassified reactions, EC classes missing assigned protein sequences, and the possibility of neglecting promiscuous candidate enzymes belonging to other EC classes. Our studies demonstrate that enrichment with functionally close promiscuous enzymes improves the functional enzyme sequence annotation compared to DeepEC and PRIAM. We also compare our method to Selenzyme^15^, a reference tool for predicting protein sequences of orphan biochemical reactions, and obtain improved predictions. We validate BridgIT+ predictions through three studies involving experimentally confirmed reaction-protein associations. Finally, we illustrate its applicability through two studies by (i) annotating novel reactions in metabolic pathway design; and (ii) proposing function for 144 poorly annotated sequences in the *E. coli* genome.

## 1.3 Results and discussion

### 1.3.1 BridgIT+ method

Briefly, the BridgIT+ workflow can perform two annotation tasks that link: (i) an orphan and novel computationally predicted reactions with proteins and (ii) orphan proteins with enzymatic functions. The core building block in both tasks is the creation of BridgIT+ PSSM profiles (Figure 1a). The input to this block is a collection of EC numbers. Whereas in our studies we select EC numbers based on promiscuity using BridgIT^10^, this input can be provided from other sources such as experimental studies and other computational prediction methods. The creation of these profiles is organized into three main steps: (1) identifying protein sequences from the UniProt database^16^ that correspond to the collected EC numbers; (2) sequence alignment using MAFFT^17^; and (3) creation of the enzymatic profiles corresponding to the aligned sequences, BridgIT+ PSSM profiles, using PSI-BLAST^18^.

**Figure 1.**
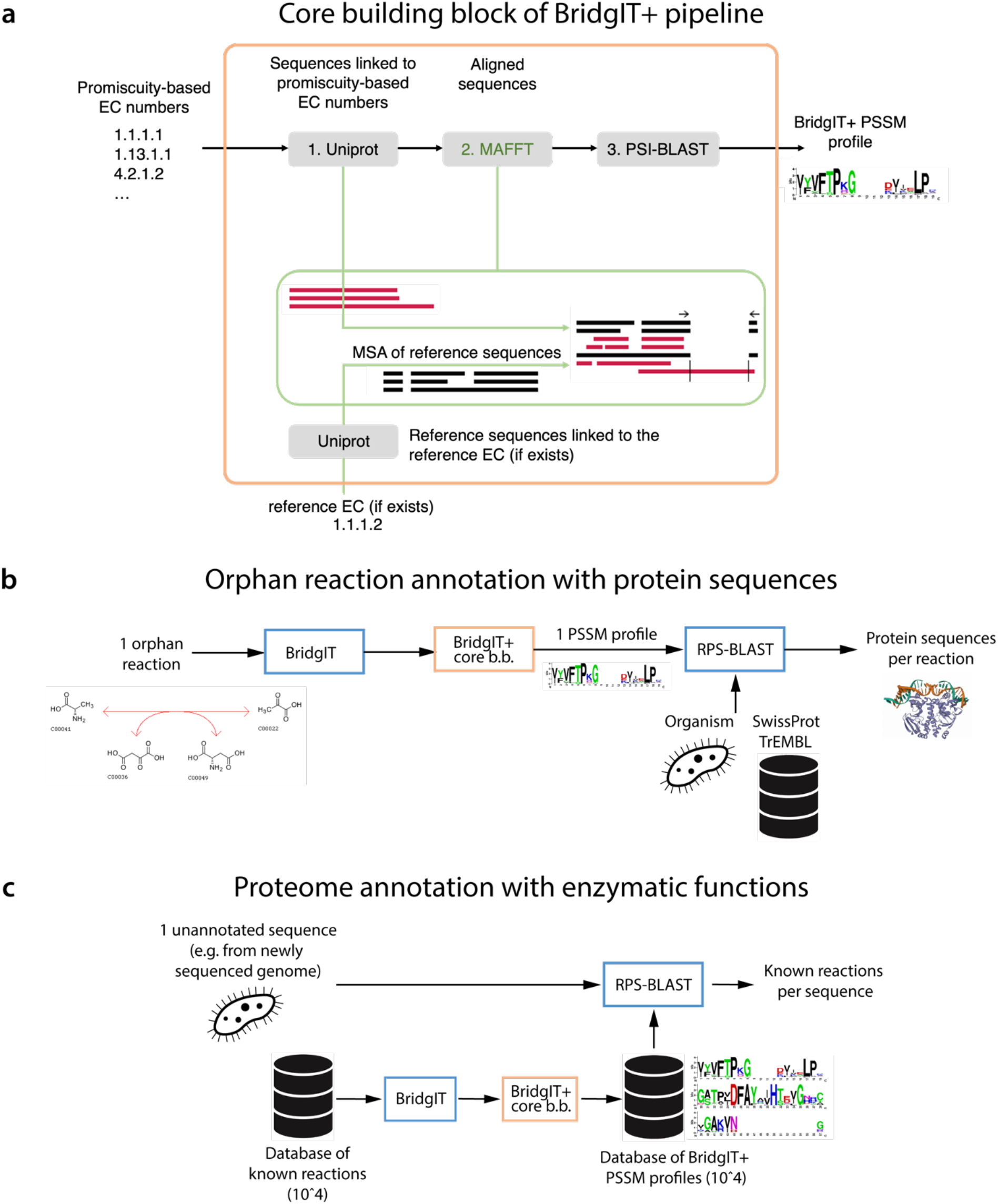
The BridgIT+ framework. **a**. General pipeline for the generation of BridgIT+ PSSM profiles structured in three steps: (1) identifying protein sequences of promiscuous enzymes (UniProt), (2) sequence alignment (MAFFT), and (3) creation of the enzymatic profiles of the aligned sequences (PSI-BLAST). BridgIT+ enzymatic profiles are employed for the annotation of: **b**. orphan reactions with enzymatic sequence and **c**. unannotated sequences with known reactions.

For annotating an orphan or a novel reaction with proteins (task (i)), the workflow uses BridgIT to provide a collection of promiscuous EC numbers corresponding to this reaction (Figure 1b). We then use this collection to compute PSSM profiles, and we screen these profiles against databases of known sequences such as TrEMBL and Swiss-Prot^19^ using RPS-BLAST^18^. The pipeline output is a ranked protein sequence set that matches the orphan or novel reaction.

To perform task (ii), annotating orphan proteins with enzymatic functions, we use RPS-BLAST to screen the orphan sequence against the previously created database of BridgIT+ PSSM profiles of known reactions (Methods). The database of BridgIT+ PSSM profiles can be extended by creating PSSM profiles of new reactions using BridgIT and the BridgIT+ core building block (Figures 1a and 1c).

### 1.3.2 Validation against biochemical assays

To assess BridgIT+ performance using experimentally confirmed reaction-protein associations, we performed three validation studies using biochemical knowledge originating from the previous versions of the KEGG database, namely v.2011 and v.2015. Using the biochemical information from the earlier versions of the KEGG database allows us to demonstrate BridgIT+ capability to predict reaction-protein relations confirmed by the later database versions (v.2021). In the validation studies, we endeavored to (A) match the orphan reactions and orphan proteins from the KEGG v.2011 database to each other, (B) use the knowledge of KEGG v.2011 protein profiles to successfully match orphan reactions to proteins later added in the KEGG database, and (C) find protein sequences for confirmed novel reactions predicted in 2015 and whose protein sequences were annotated between 2015 and 2021 (**Figure 2**).

**Figure 2.**
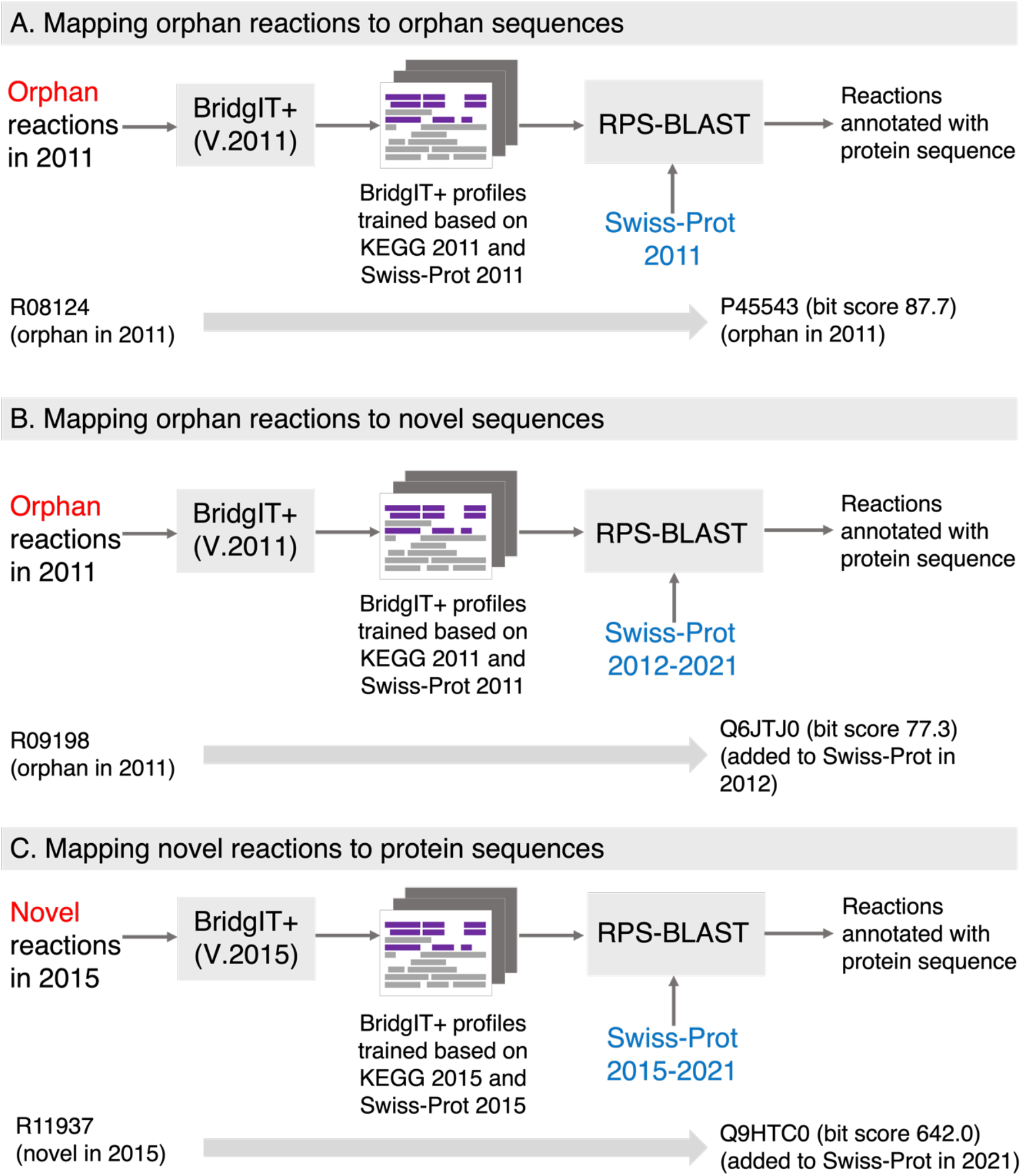
Design of validation studies of BridgIT+ against biochemical assays. **a**. Mapping orphan reactions to orphan sequences, both available in public databases in 2011, but not linked to each other. **b**. Mapping orphan reactions from 2011 to novel sequences added to public databases in 2012-2021. **c**. Mapping novel reactions computationally predicted in 2015 to protein sequences added to public databases in 2015-2021.

### A. BridgIT+ correctly matches orphan reactions to orphan sequences

In the KEGG and UniProt databases released in 2011, we found seven orphan reactions that, in the later versions of KEGG, became associated with orphan protein sequences of UniProt v.2011. To be more specific, the UniProt v.2011 already contained protein sequences capable of catalyzing these reactions, but their functions still needed to be discovered. Their enzymatic activities were later discovered, assigned to new EC numbers, and linked to the KEGG reactions. Matching directly orphan reactions to orphan sequences is more challenging than assigning a reaction to an EC number and an EC number to a protein sequence because the EC numbers corresponding to these enzymatic activities were unknown in 2011. Indeed, methods that require information about the EC classification, such as DeepEC and PRIAM, cannot be used to perform this task. Since these enzymatic activities have been experimentally confirmed using biochemical assays, we used these seven reactions as a benchmark to evaluate the BridgIT+ performance.

To this end, we formed the BridgIT reference reaction database using the reactions from KEGG v.2011, and we created the BridgIT+ protein profiles based on the protein sequences in Swiss-Prot v.2011 (Methods). The BridgIT+ profiles represent the alignment of promiscuous protein sequences proposed by BridgIT based on the structural similarity of their reactions. We then performed a homology search between each orphan protein in UniProt v.2011 and BridgIT+ profiles using the RPS-BLAST program. Finally, we compared the BridgIT+ annotation results with the approved enzyme assignments in later versions of KEGG. Remarkably, BridgIT+ matched the seven orphan reactions to their correct orphan sequences and three-level EC numbers corresponding to the reaction mechanism (**Table 1**).

**Table 1.**
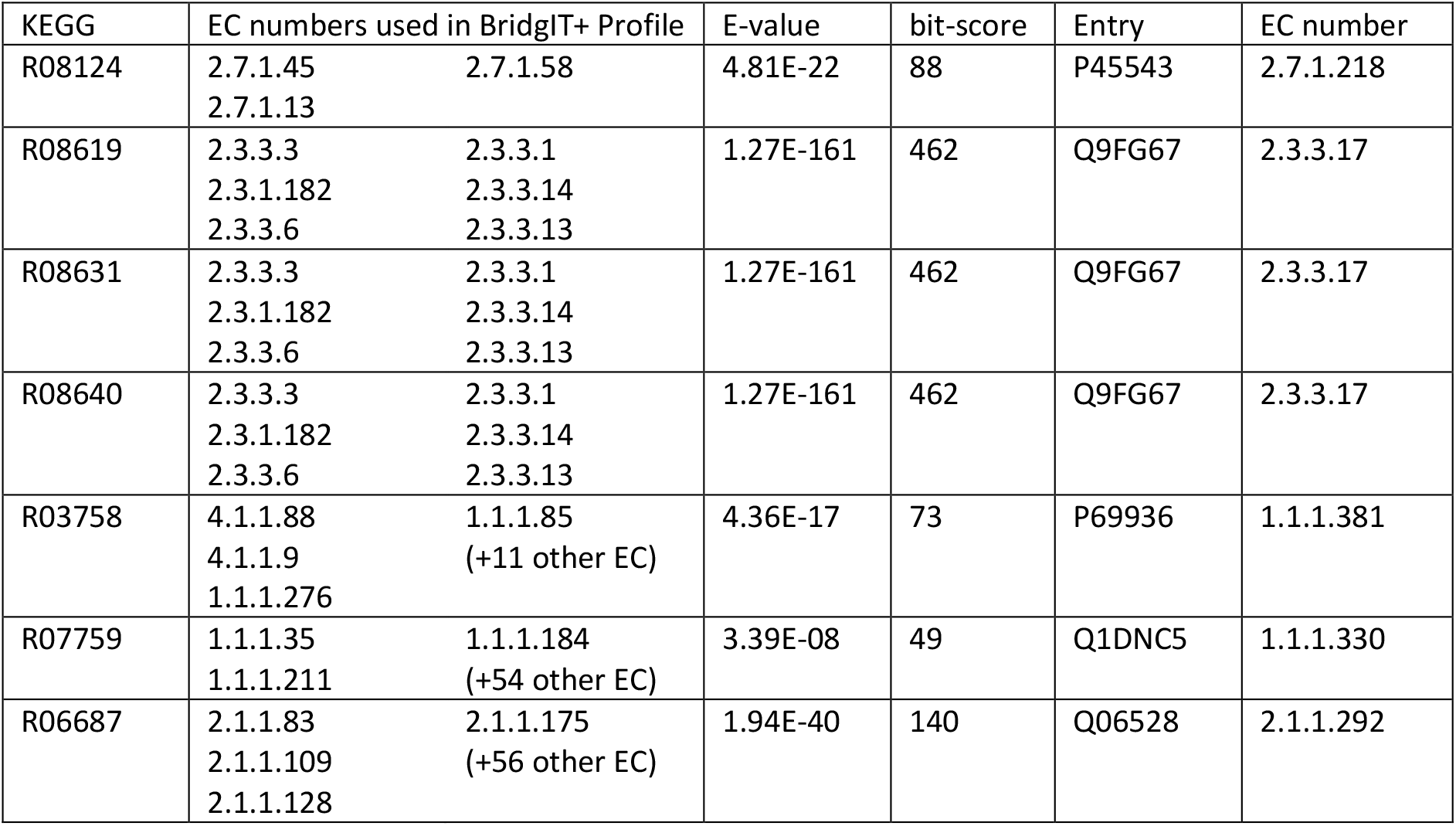
Annotation of formerly orphan KEGG reactions with protein sequence using the 2011 version of BridgIT+ reference profiles.

### B. BridgIT+ correctly matches orphan reactions to newly added protein sequences

We used the trained profiles based on information from 2011 to determine whether BridgIT+ can link the orphan reactions in 2011 to the correct protein sequences added between 2012 to 2021. Out of 234 orphan reactions in KEGG 2011 that became later non-orphan, 75 were structurally balanced and assigned to a new EC number, and we used them to test BridgIT+. 37 out of 75 reactions were correctly mapped to their protein sequences with the exact four-level EC number, demonstrating BridgIT+ ability to match reactions to enzymes with correct substrate specificity (Supplementary Table 1, bit-score: 23 to 1084). More strikingly, all 75 reactions were linked to a protein sequence with the correct three-level EC number, indicating that our method matches reactions to enzymes with correct reaction mechanisms. In other words, BridgIT+ trained only on biochemical information from 2011 can correctly assign the orphan reactions from 2011 to protein sequences confirmed in later years.

### C. BridgIT+ correctly matches novel reactions to protein sequences

The ATLAS of Biochemistry databases^20–23^ provide a comprehensive source of theoretically possible biochemical reactions. The first version of this database was centered around KEGG compounds available in 2015. In 2020, we found that the newly available biochemical data validated 107 novel reactions predicted in ATLAS 2015^21^. Here, we examined the capability of BridgIT+ to assign correct protein sequences to these novel reactions. From 2015, Swiss-Prot sequences were assigned to 83 out of 107 formerly novel reactions according to KEGG and Rhea databases. For these 83 reactions, we compared the validated protein sequence annotation with BridgIT+ predictions. These reactions have up to 255 unique validated sequences, with a median of 4 sequences per reaction. For 70 reactions (84%), experimentally assigned sequences identically matched the Swiss-Prot identifiers predicted by BridgIT+ (Supplementary Table 2). For 9 out of the remaining 13 reactions the BridgIT+ top-ranked enzyme sequence corresponded to the reaction mechanism with the identical three-level EC number as the queried reaction (Supplementary material 2). For one of the remaining 13 reactions, only one protein sequence was available for the EC classes predicted with BridgIT; since this sequence was correctly matched, the BridgIT+ pipeline was superfluous. For the other two out of the remaining 13 reactions, even though the best EC number prediction according to BridgIT was correct, BridgIT+ could correctly capture the reaction mechanism only if we neglected the enzyme promiscuity (Supplementary material). Finally, one reaction did not have information about the reaction mechanism because only a two-level EC number was assigned to it (Supplementary material). Overall, these results demonstrate the predictive capabilities of BridgIT+ for novel reactions because it assigned identical or matching rection mechanism protein sequences for 79 of 83 formerly novel reactions (95%).

**Table 2.**
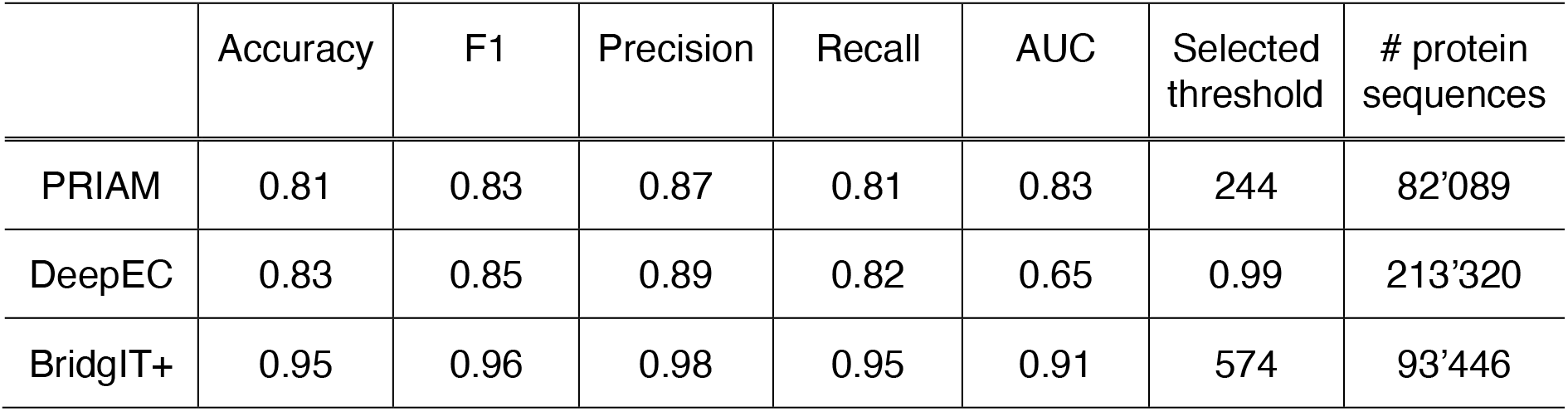
Comparison of the accuracy, F1 score (the harmonic mean of precision and recall), precision, recall, and AUC (the area under the curve) for the three tools annotating Swiss-Prot sequences. The selected threshold (bit-score for PRIAM and BridgIT+, the score for DeepEC) is based on the best F1 score. The number of processable by the tool unique protein sequences is indicated for each tool.

We next compared the predictions for KEGG and Rhea databases separately. Of 60 reactions with manually annotated protein sequences in Rhea, 51 had a correct protein sequence assigned with BridgIT+. Similarly, for 50 out of 59 reactions with Swiss-Prot protein annotation in KEGG, BridgIT+ performed correct sequences assignment. These results indicate that BridgIT+ is agnostic to the source of reactions and that it performs equally well for the Rhea and KEGG databases.

### 1.3.3 Comparison with other tools

#### Protein sequence annotation

Annotating sequences with enzymatic function is valuable for the prediction of metabolic phenotypes of organisms based on sequenced genomes. We compared the protein annotation performance of BridgIT+ (using the EC-based profiles) with the representative tools in the field: DeepEC and PRIAM.

We evaluated the prediction performances of the three tools for annotating Swiss-Prot sequences (Table 2). We selected Swiss-Prot as ground truth for comparison because it has been expertly annotated using a state-of-the-art methodology (automated screening) and current biological knowledge (human inspection). For each enzyme sequence, we counted how many enzyme activities predicted by these tools were reported in Swiss-Prot (true positive, TP), not reported (probable false positive, FP), and how many activities reported in Swiss-Prot were missed by the tool (false negative, FN). We evaluated the quality of the predictions using the accuracy 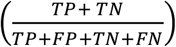, precision 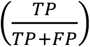,recall 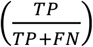, F1 score 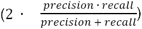, and area under the curve (AUC) measures. Since there is no systematic testing for the absent enzymatic activities of enzymes reported to Swiss-Prot, the true negative (*TN*) measure and associated statistical metrics could not be estimated.

BridgIT+ outperformed PRIAM and DeepEC in this task regarding all measured performance indicators (Table 2). We obtained a larger area under the ROC^24^ curve (AUC) in BridgIT+ predictions (0.91) compared to PRIAM (0.83) and DeepEC (0.65). Moreover, BridgIT+ precision and recall indicators (0.98 and 0.95, respectively) were superior compared to DeepEC (0.89, 0.82) and PRIAM (0.87, 0.81). We also noted that BridgIT and PRIAM demonstrate a better ability to discriminate the EC numbers than DeepEC due to the advantage of homology-based tools over the neural networks for the sequence annotation with few known homology instances per EC number. We argue that the improvement in the performance of BridgIT+ was achieved by enriching the profiles with promiscuous sequences assigned to alternative EC numbers.

#### Reaction annotation with protein sequences

To comparatively evaluate BridgIT+ performance in this task, we used the standard tool for linking reactions to protein sequences, Selenzyme^15^. Selenzyme was published in 2018, and we could not modify the tool’s reference set, train it based on information from 2011, and perform a fair comparison. Nevertheless, we used the current version of Selenzyme to annotate the seven orphan reactions that BridgIT+ correctly matched to the orphan sequences (see *Validation against biochemical assays* and Supplementary Table 3). In the list of top 50 candidates per reaction Selenzyme provided, we could find a protein sequence with the correct EC number for only 5 out of 7 reactions. This result suggests that BridgIT+ performs better than Selenzyme for establishing protein-reaction associations.

### 1.3.4 Applications

#### BridgIT+ facilitates enzyme discovery in metabolic pathway design

Microbial biosynthesis is one of the most effective approaches to producing complex compounds such as natural pharmaceuticals^25^. To guide and accelerate the design of biosynthesis pathways toward biochemicals, computational tools for enzyme discovery are essential. In a recent study, Srinivasan et al. implemented the conversion of hyoscyamine and scopolamine to cognate *N*-oxides to produce natural plant products in yeast using two novel ATLAS reactions along with their putative enzyme candidates suggested by BridgIT^26^. BridgIT proposed senecionine *N*-oxygenase (EC 1.14.13.101) as the best candidate for both conversions. Srinivasan et al. analyzed the activity of 3 orthologs of this enzyme from three different species in yeast: *Tj*SNO from *Tyria jacobaeae* (cinnabar moth), *Gg*PNO from *Grammia geneura* (Nevada tiger moth), and *Zv*PNO from *Zonocerus variegatus* (harlequin locust). They finally reported the highest heterologous production of hyoscyamine *N*-oxide and scopolamine *N*-oxide by the yeast strain expressing *Zv*PNO.

Here, instead of proposing EC numbers, we go further with BridgIT+ and annotate these two novel ATLAS reactions with a ranked list of candidate protein sequences. If BridgIT+ were available at the time, it would replace the manual work of selecting the best protein candidates for the EC numbers. Following the BridgIT+ pipeline, we trained an enzyme promiscuity-enriched profile for each novel reaction. Then, we used RPS-BLAST to search and rank the enzyme orthologs from different species in Swiss-Prot and TrEMBL databases (Supplementary Table 4). Finally, we compared the results of BridgIT+ with the experimental results by Srinivasan et al. In the ranking results of BridgIT+, *Zv*PNO (bit scores: 96.8 and 94.5) ranked higher compared to *Tj*SNO (with bit scores 76.4, 74) and *Gg*PNO (with bit scores: 74, 71.3), thus matching the experimental observations closely. These results indicate that BridgIT+ can provide precise enzyme sequence predictions for annotating computationally predicted reactions, facilitating the metabolic pathway design.

#### BridgIT+ correctly annotated enzyme activities in the whole genome of *E. coli* and proposed function for 144 poorly annotated sequences

The rate of protein functional elucidation needs to catch up to the pace of gene and protein sequence discovery, leading to an accumulation of proteins with unknown functions. *Escherichia coli*, perhaps the best-studied model organism extensively annotated in the Swiss-Prot database, is not an exception. Recent studies show that 1’431 proteins (35%) of *E. coli* are still not functionally annotated^27^.

To analyze the capabilities of BridgIT+ for genome annotation, we applied BridgIT+ profiles to the *E. coli* K-12 proteome and compared BridgIT+ results with the manual annotation in Swiss-Prot (**Figure 3**, A). Out of 1’130 EC numbers linked to 6’066 protein entries in Swiss-Prot, 603 EC numbers associated with 686 proteins could be processed by BridgIT+ (Methods). Of the 603 EC numbers, 598 (98%) linked to 660 protein sequences were correctly annotated by BridgIT+ (**Figure 3**, B). The remaining five EC numbers had assigned highly promiscuous EC numbers with very broad substrate specificity (3.5.2.6, 3.1.1.5, 2.5.1.18, 3.1.3.2, and 3.2.1.21). A possible way to annotate enzymes with such broad substrate specificity with BridgIT+ is to remove the promiscuity from consideration. This way, the BridgIT+ profile would be more specific to substrates of metabolic reactions during the homology search.

**Figure 3.**
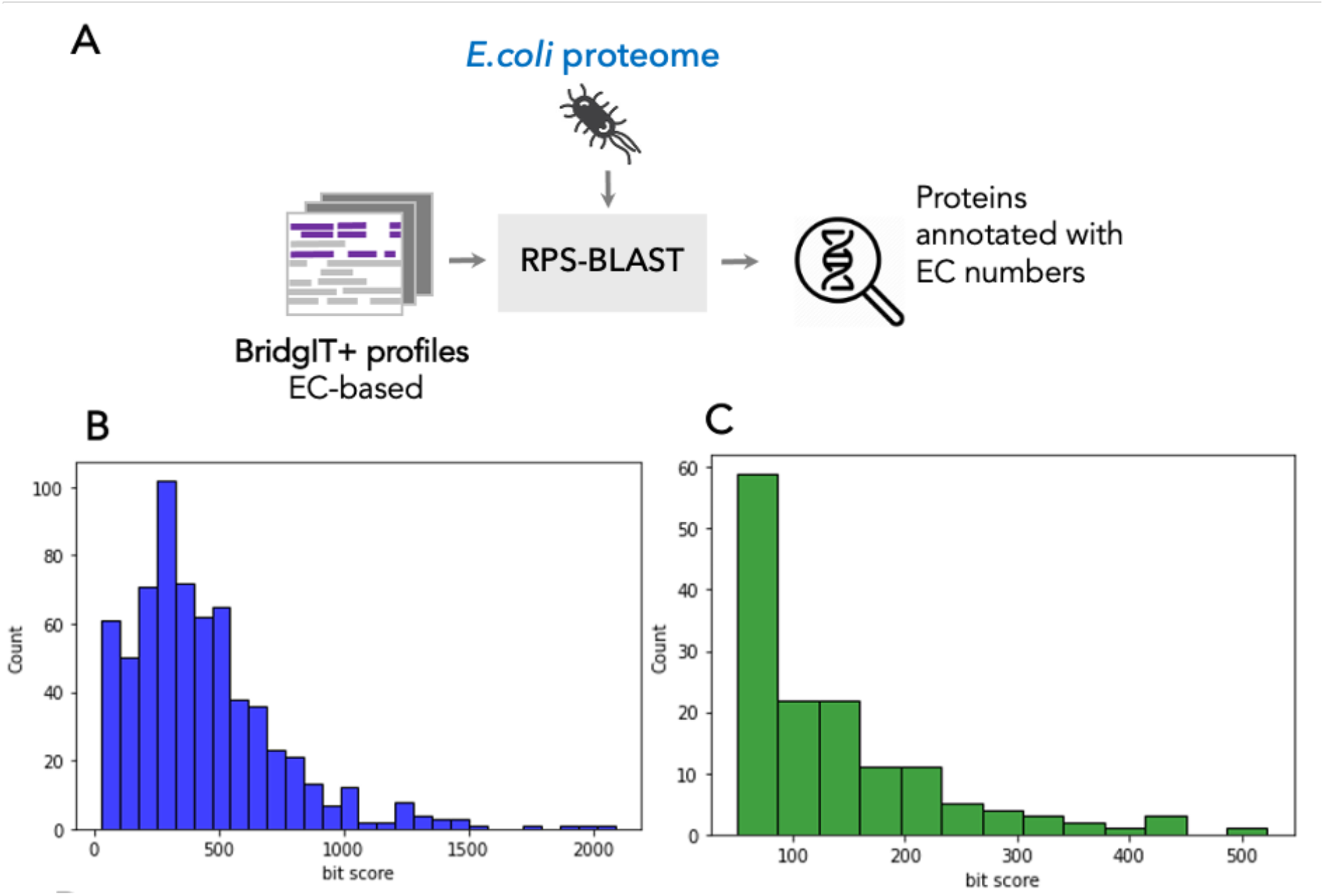
Annotation of *E. coli* proteome using BridgIT+ reference profiles. **a**. Schematic representation of the approach used to annotate an organism proteome with BridgIT+ profiles. **b**. Bit-score for 598 reactions correctly linked to 660 protein sequences by BridgIT+. **c**. Bit-score for 144 *E. coli* sequences without 4-level EC number annotation assigned with 110 unique four-level EC numbers using BridgIT+ profiles.

We next used BridgIT+ to address the need for genome functional annotation. BridgIT+ annotated 144 *E. coli* sequences currently missing a four-level EC number with 110 unique four-level EC numbers (Figure 3, C). The identified EC numbers correspond to reactions not currently cataloged in the *E. coli* metabolism. These results suggest that BridgIT+ can fill the metabolic gaps and annotate new biochemical pathways in any sequenced genome.

## 1.4 Conclusions

The wealth of genome and proteome data calls for robust methods for the functional annotation of genes and proteins. A prominent approach leveraging multiple sequence alignment for annotations is PRIAM^4^. It collects sequences per EC number and creates position-specific scoring matrices (PSSM) for homologous modules found within each enzyme-specific collection. Recently, a deep learning approach DeepEC^6^ outperformed PRIAM primarily due to its sensitivity to nonlinearities introduced by mutated domains and binding site residues. Still, scarce data for some EC classes hinder DeepEC’s performance, a common drawback of deep learning approaches. Both DeepEC and PRIAM link enzyme functionality to protein sequences through EC numbers. However, knowledge of EC numbers is not always sufficient for establishing protein-reaction associations because multiple proteins can catalyze reactions with a single four-level EC number. Additionally, EC numbers do not capture the evolutionary changes of enzymes. Indeed, it was reported that approximately 40% of enzymes have evolved to completely new functionality, i.e., to EC classes differing in the first digit of EC^9^. Therefore, relying entirely on EC can lead to erroneous predictions.

Here proposed method, BridgIT+, bases its reaction-protein associations on the individual reaction mechanisms rather than on a human-error-prone EC classification. In other words, whereas EC-based approaches indicate which proteins best represent EC numbers, BridgIT+ answers which proteins best represent reactions. Similar to the previously proposed method BridgIT, which outperformed peers for reaction annotation with EC numbers^16^, BridgIT+ leverages knowledge of reaction mechanisms and enzyme promiscuity to annotate reactions with protein sequences likely to catalyze these reactions. BridgIT+ can be used on current reaction and protein databases to draw meaningful functional associations between the two. Building this direct link allows us to go beyond EC numbers and provide interpretable predictions for sequences and reactions lacking EC annotation.

Comparing our method to experimental biochemical assays, we demonstrated that BridgIT+ could successfully match the orphan reactions and orphan proteins available in the databases but missed the link to each other. We have also shown that our method can successfully use the current biochemical knowledge contained in reaction and protein databases to match future orphan reactions and proteins.

BridgIT+ brings significant advantages compared to existing methods as it does not require extensive data, e.g., for neural network training, and adds promiscuity into consideration to improve the prediction in case of the absence of homology. Compared to DeepEC, PRIAM, and Selenzyme, the observed improvement in performance is brought about by grouping reference sequences based on their catalytic function. BridgIT+ produces a ranked list of protein sequences for any reaction and can be adapted to any specific organism or application. We expect that BridgIT+ predictive capabilities will grow as more enzyme sequences are introduced into protein sequence databases and more reaction mechanisms are discovered and cataloged.

## 1.5 Methods

### Curating input for BridgIT+

The input for BridgIT+ is generated using BridgIT^10^. The standard BridgIT output has a similarity score for each predicted EC number. The reactive site is identified using BNICE.ch^13^ reaction rules, the fingerprint is generated, and the EC number and score are predicted as described in the original BridgIT publication. Alternatively, any set of EC numbers per reaction can be used as input.

### Constructing EC-specific profiles with BridgIT+

As input to the BridgIT+ workflow for creating profiles, we collected known EC numbers (reference ECs). To create a BridgIT+ profile, the collected ECs (1) needed to have a four-level EC number defined, (2) were linked to at least two protein sequences in total, (3) were linked to at least one metabolic reaction in the public databases, (4) the linked reaction was reconstructed with an enzymatic rule, and (5) the linked reaction participants were fully structured (e.g., not including proteins or polymers). ECs satisfying criteria (1)-(5) were eligible for BridgIT+ processing and were considered for proteome annotation. Similarly, orphan reactions for which a profile is created should be reconstructed with an enzymatic reaction rule and be fully structured to be processable. The reference EC numbers were used to query the LCSB database to find all linked biochemical reactions. Next, we used BridgIT to find the most similar reactions to the extracted biochemical reactions with the reactive site-centric fingerprints. The EC classes were collected from the BridgIT output depending on the prediction score threshold and distance from the reactive site (level). The EC numbers associated with the most similar reactions designated the candidate’s promiscuous activities. The ranked list of EC numbers was used to collect sequences from protein databases (such as UniProt^16^). We used the MAFFT (Multiple Sequence Alignment by the Fast Fourier Transform) method^17^ to align reference sequences with clustered promiscuous sequences. MAFFT method begins by creating a Multiple Sequence Alignment (MSA) of the reference sequences, then aligns the promiscuous sequences cluster to the reference MSA (joint MSA). Joint MSA preserves the biochemical knowledge of the reference EC number and takes promiscuity into account. Finally, the alignment was used to generate enzymatic profiles (BridgIT+ profiles) using PSI-BLAST.

### Predicting protein sequence for an orphan reaction

A single orphan reaction could be an input for the BridgIT+ workflow to construct an orphan reaction-specific profile and predict a protein to catalyze it. First, BridgIT was used to find the most similar reactions with complete EC class annotation. Corresponding EC classes were collected, linked sequences requested from UniProt, aligned, and used to construct the BridgIT+ profile for the orphan reaction. This profile can be used to find a potential catalyzing sequence within a specific organism or the whole set of sequences using RPS-BLAST.

### Sequence prediction and ranking for each BridgIT+ level profile

A collection of reference EC numbers can be used as BridgIT+ input for protein function annotation. After creating BridgIT profiles for all the reference EC numbers, they can be used for annotating a genome using RPS-BLAST. Matching a BridgIT+ profile to a sequence implies the catalytic activity and the corresponding reference EC number, with a bit-score of more than 50 signifying the confidence of the predicted EC annotation.

### Comparison of BridgIT+ performance to PRIAM and DeepEC

In comparing BridgIT+ with related tools, we have used a standard procedure and the optimal thresholds for each tool. The following optimal thresholds were used: a bit score of 50 for BridgIT+, e-value of 10^-30^ for PRIAM, and default hyperparameters of DeepEC.

### Swiss-Prot database download

The set of annotated protein sequences was downloaded from https://www.uniprot.org/uniprotkb?query=reviewed:true in August 2018.

### Acquiring Selenzyme results

Selenzyme results were retrieved in December 2021 based on KEGG identifiers from http://selenzyme.synbiochem.co.uk.

## 1.6 Data and code availability

The data and scripts used to produce, analyze, and visualize the results are available at https://doi.org/10.5281/zenodo.8268529. The code for the BridgIT+ pipeline is available at https://github.com/EPFL-LCSB/BridgITplus.

## 1.7 Acknowledgments

Funding for this work was provided by the Swiss National Science Foundation (SNSF grant agreement 200021_188623 and NCCR Microbiomes grant agreement 51NF40_180575), the European Union’s Horizon 2020 research and innovation program under grant agreement 814408 and Marie Skłodowska-Curie grant agreement No 72228, and the Ecole Polytechnique Fédérale de Lausanne (EPFL).

## References

1. Baric, R. S., Crosson, S., Damania, B., Miller, S. I. & Rubin, E. J. Next-generation highthroughput functional annotation of microbial genomes. mBio 7, (2016).

2. Griesemer, M., Kimbrel, J. A., Zhou, C. E., Navid, A. & D’Haeseleer, P. Combining multiple functional annotation tools increases coverage of metabolic annotation. BMC Genomics 19, (2018).

3. Sinha, S., Lynn, A. M. & Desai, D. K. Implementation of homology based and non-homology based computational methods for the identification and annotation of orphan enzymes: using Mycobacterium tuberculosis H37Rv as a case study. BMC Bioinformatics 21, (2020).

4. Claudel-Renard, C., Chevalet, C., Faraut, T. & Kahn, D. Enzyme-specific profiles for genome annotation: PRIAM. Nucleic Acids Res 31, 6633–6639 (2003).

5. Bileschi, M. L. et al. Using deep learning to annotate the protein universe. Nat Biotechnol 40, 932–937 (2022).

6. Ryu, J. Y., Kim, H. U. & Lee, S. Y. Deep learning enables high-quality and highthroughput prediction of enzyme commission numbers. Proc Natl Acad Sci U S A 116, 13996–14001 (2019).

7. Vamathevan, J. et al. Applications of machine learning in drug discovery and development. Nature Reviews Drug Discovery vol. 18 463–477 Preprint at 10.1038/s41573-019-0024-5 (2019).

8. Pearson, W. R. An introduction to sequence similarity (‘homology’) searching. Curr Protoc Bioinformatics (2013) doi:10.1002/0471250953.bi0301s42.

9. Martínez Cuesta, S., Rahman, S. A., Furnham, N. & Thornton, J. M. The Classification and Evolution of Enzyme Function. Biophysical Journal vol. 109 1082–1086 Preprint at 10.1016/j.bpj.2015.04.020 (2015).

10. Hadadi, N., MohammadiPeyhani, H., Miskovic, L., Seijo, M. & Hatzimanikatis, V. Enzyme annotation for orphan and novel reactions using knowledge of substrate reactive sites. Proc Natl Acad Sci U S A 116, 7298–7307 (2019).

11. Hadadi, N. & Hatzimanikatis, V. Design of computational retrobiosynthesis tools for the design of de novo synthetic pathways. Curr Opin Chem Biol 28, 99–104 (2015).

12. Sveshnikova, A., MohammadiPeyhani, H. & Hatzimanikatis, V. Computational tools and resources for designing new pathways to small molecules. Curr Opin Biotechnol 76, 102722 (2022).

13. Hatzimanikatis, V. et al. Exploring the diversity of complex metabolic networks. Bioinformatics 21, 1603–1609 (2005).

14. Tokić, M. et al. Discovery and Evaluation of Biosynthetic Pathways for the Production of Five Methyl Ethyl Ketone Precursors. ACS Synth Biol acssynbio.8b00049 (2018) doi:10.1021/acssynbio.8b00049.

15. Carbonell, P. et al. Selenzyme: Enzyme selection tool for pathway design. Bioinformatics 34, 2153–2154 (2018).

16. Bateman, A. et al. UniProt: the universal protein knowledgebase in 2021. Nucleic Acids Res 49, D480–D489 (2021).

17. Katoh, K., Rozewicki, J. & Yamada, K. D. MAFFT online service: Multiple sequence alignment, interactive sequence choice and visualization. Brief Bioinform 20, 1160–1166 (2018).

18. Sayers, E. W. et al. Database resources of the National Center for Biotechnology Information in 2023. Nucleic Acids Res 51, D29–D38 (2023).

19. Bairoch, A. & Apweiler, R. The SWISS-PROT protein sequence database and its supplement TrEMBL in 2000. Nucleic Acids Res 28, 45–48 (2000).

20. Hadadi, N., Hafner, J., Shajkofci, A., Zisaki, A. & Hatzimanikatis, V. ATLAS of Biochemistry: A Repository of All Possible Biochemical Reactions for Synthetic Biology and Metabolic Engineering Studies. ACS Synth Biol 5, 1155–1166 (2016).

21. Hafner, J., MohammadiPeyhani, H., Sveshnikova, A., Scheidegger, A. & Hatzimanikatis, V. Updated ATLAS of Biochemistry with New Metabolites and Improved Enzyme Prediction Power. ACS Synth Biol 9, 1479–1482 (2020).

22. MohammadiPeyhani, H., Hafner, J., Sveshnikova, A., Viterbo, V. & Hatzimanikatis, V. Expanding biochemical knowledge and illuminating metabolic dark matter with ATLASx. Nat Commun 13, 1–12 (2022).

23. Sveshnikova, A., MohammadiPeyhani, H. & Hatzimanikatis, V. ARBRE: Computational resource to predict pathways towards industrially important aromatic compounds. Metab Eng 72, 259–274 (2022).

24. McNeil, B. J. & Hanley, J. A. Statistical Approaches to the Analysis of Receiver Operating Characteristic (ROC) Curves. Medical Decision Making 4, 137–150 (1984).

25. Pham, J. v. et al. A review of the microbial production of bioactive natural products and biologics. Front Microbiol 10, (2019).

26. Srinivasan, P. & Smolke, C. D. Engineering cellular metabolite transport for biosynthesis of computationally predicted tropane alkaloid derivatives in yeast. Proc Natl Acad Sci U S A 118, (2021).

27. Ghatak, S., King, Z. A., Sastry, A. & Palsson, B. O. The y-ome defines the 35% of Escherichia coli genes that lack experimental evidence of function. Nucleic Acids Res 47, 2446–2454 (2019).

