## Supplementary material 2 for "Enzyme promiscuous profiles for protein sequence and reaction annotation"

| **Reaction** | **Externally assigned Swiss-Prot ID** | **Externally assigned EC numbers** | **BridgIT+ assigned Swiss-Prot ID (bit-score > 50)** | **Best sequence alternative according to BridgIT+** | **Distance from reactive site used*** |
| --- | --- | --- | --- | --- | --- |
| R08480 | P0DPA8 | 2.7.1.222 | None | None | L1 |
| R11785 | Q87BG6,Q4UVL3,Q9PAM6,Q9HZ62,Q8P8H3,Q88M11 | 3.1.3.105 | None | None | L0 |
| R10964 | B0CN31 | 2.1.1.304 | None | None | L3 |
| R11772 | Q9WYP7 | 5.1.-.- | None | None | L7 |
| R11358 | A0R4Q6,P9WPL4,P9WPL5,P9WPP3 | 1.14.15.28 | 47 | Q4QQV7  (bit-score: 290) | L7 |
| R11552 | Q8IY26,Q5TZ07,Q66H88,Q9D4F2,Q58DI5 | No | 15 | Q01968  (bit-score: 590) | L1 |
| R11553 | Q9UKK9,Q5RCY2,Q6AY63,Q9JKX6 | 2.7.7.96 | 315 | Q8NFF5  (bit-score: 812) | L5 |
| R11726 | Q6LYX3,O34895,Q8PYC8 | 1.1.1.409, 2.3.1.264 | 335 | P58718  (bit-score 366) | L2 |
| R11901 | A7IPX7 | 1.14.13.69 | 10 | O87082  (bit-score: 730) | L1 |
| R11333 | B9JN19,Q6D8V9,A0QXE5,A0R757,Q9ZB26 | 5.3.1.34 | 393 | B7MM49  (bit-score 273) | L2 |
| R11313 | Q9K9G9,O31724,Q9ASY9,Q9SWB6,B6TPF2 | 3.5.1.- | 9 | Q99497  (bit-score: 157) | L3 |
| R11145 | Q9BTZ2,Q5RCF8,A7B3K3 | 1.1.1.392 | 516 | Q3ZC42  (bit-score 312) | L7 |
| R09055 | Q02430,D5AP79 | 1.1.1.396 | 1174 | G5DGA8  (bit-score: 195) | L3 |

* L0 – only the atoms of reactive site, no bonds

L1 – atoms and bonds of reactive site

L2 – reactive site + 1 atom away from the reactive site

L3 – reactive site + 2 atoms away from the reactive site

… and so on until L7

**Possible reasons for difference in external assignment and BridgIT+ predictions**

**Too much promiscuity**

If the set of ECs does not correspond to the shared reaction mechanism, then the PSSM profile does not reflect the reaction mechanism and therefore no reliable predictions below the e-value cut-off of 10 could be found. Too much promiscuity usually appears on the lower levels of BridgIT predictions.

For reactions R08480 and R11785 no promiscuous predictions available on the levels above L1 (used for R08480) and L0 (used for R11785). The BridgIT predictions comprised only 1 correct EC number (2.7.1.222 and 3.1.3.105) on the levels above L1 and L0.

When only L7 (without promiscuous results for these two cases) was considered, correct BridgIT+ results were obtained:

| **KEGG**  **ID** | **Expected SwissProt**  **ID** | **Predicted sequence** | **BridgIT+ profile** | **%identity** | **alignment length** | **mis-**  **matches** | **evalue** | **bit score** |
| --- | --- | --- | --- | --- | --- | --- | --- | --- |
| R08480 | P0DPA8 | sp\|P0DPA8\|  PSIK_PSICU | R08480L7_  A0A286LEZ6 | 73.481 | 362 | 95 | 0 | 691 |
| R11785 | Q88M11 | sp\|Q88M11\|  MUPP_PSEPK | R11785L7_  Q88M11 | 100 | 223 | 0 | 2.83E-152 | 410 |
| R11785 | Q9HZ62 | sp\|Q9HZ62\|  MUPP_PSEAE | R11785L7_  Q88M11 | 71.749 | 223 | 63 | 4.11E-151 | 407 |
| R11785 | Q8P8H3 | sp\|Q8P8H3\|  GPH_XANCP | R11785L7_  Q88M11 | 44.976 | 209 | 114 | 1.84E-60 | 177 |
| R11785 | Q4UVL3 | sp\|Q4UVL3\|  GPH_XANC8 | R11785L7_  Q88M11 | 44.976 | 209 | 114 | 1.84E-60 | 177 |
| R11785 | Q87BG6 | sp\|Q87BG6\|  GPH_XYLFT | R11785L7_  Q88M11 | 41.748 | 206 | 119 | 2.34E-57 | 170 |
| R11785 | Q9PAM6 | sp\|Q9PAM6\|  GPH_XYLFA | R11785L7_  Q88M11 | 41.262 | 206 | 120 | 1.52E-55 | 165 |
| R11785 | Q8P8H3 | sp\|Q8P8H3\|  GPH_XANCP | R11785L7_  Q88M11 | 45.455 | 11 | 6 | 6.5 | 13.9 |
| R11785 | Q4UVL3 | sp\|Q4UVL3\|  GPH_XANC8 | R11785L7_  Q88M11 | 45.455 | 11 | 6 | 6.5 | 13.9 |

**Too little promiscuity info available**

R10964: only 1 sequence (with correct EC assigned by BridgIT), no promiscuous profiles possible

**Incomplete EC + isomerization reactions**

R11772: tree of isomerization reactions obtained with correctly predicted EC 5.1.-.- was too promiscuous. Isomerization reactions are a big group. No profile that would provide predictions below the e-value cut-off of 10 were found.

**BridgIT+ predictions match well the reaction mechanism (see figures comparison).**

R11358: proposed enzyme is of EC number 1.14.99.38 instead of 1.14.15.28. Catalyzes the dehydration of cholesterol molecule as well.

R11552: proposed enzyme is of EC number 3.1.3.36. No EC assigned to the original reactions, but it is catalyzed by phosphatase as well.

R11553: proposed enzyme is of EC number 2.7.7.2 instead of 2.7.7.96. Phosphatase with the group transfer as well.

R11726: proposed enzyme is of EC number 1.1.1.408 instead of 1.1.1.409. Identical reaction mechanism, the only distinction is stereochemistry.

R11901: proposed enzyme is of EC number 1.14.13.69, same as externally assigned 1.14.13.69. Identical reaction mechanist, vinyl chloride as substrate instead of propene.

R11333: proposed enzyme is of EC number 5.3.1.17 instead of 5.3.1.34. Identical reaction mechanism, similar substrate structures

R11313: proposed enzyme is of EC number 3.5.1.124 instead of 3.5.1.-

R11145: proposed enzyme is of EC number 1.1.1.1 instead of 1.1.1.392

R09055: proposed enzyme is of EC number 1.1.1.386 instead of 1.1.1.396

**Figures comparison:**

R11358

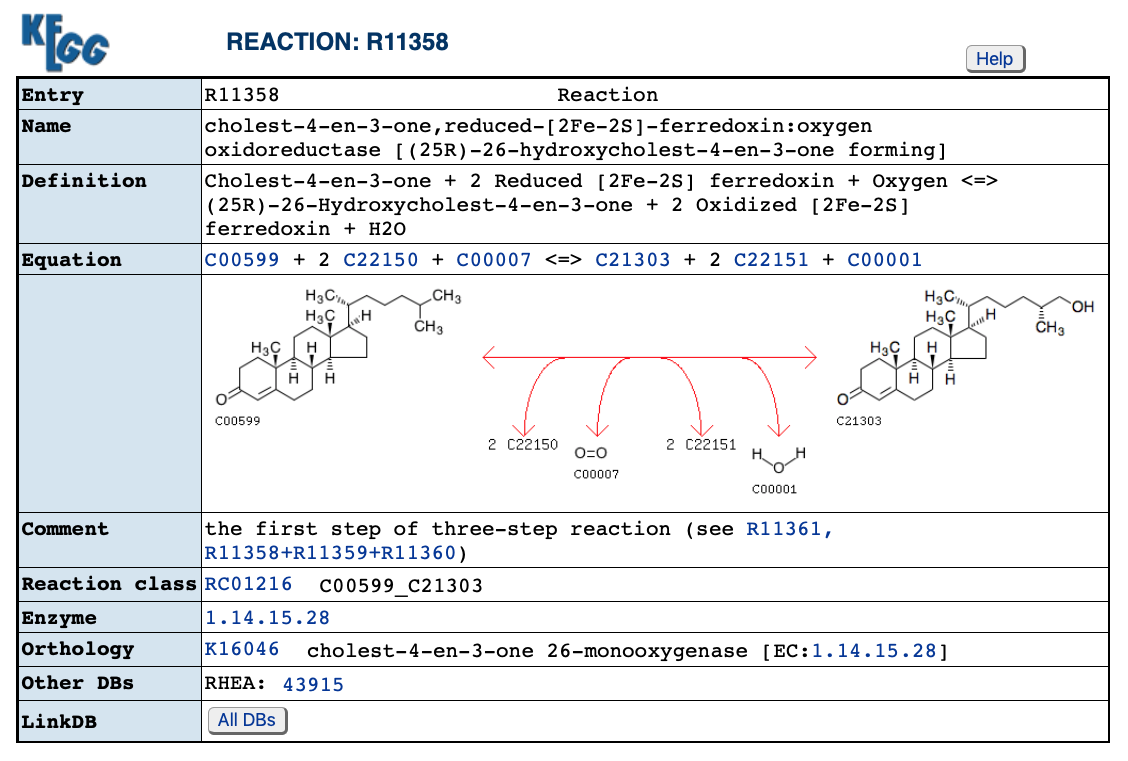

Reaction of the best protein prediction according to BridgIT+:

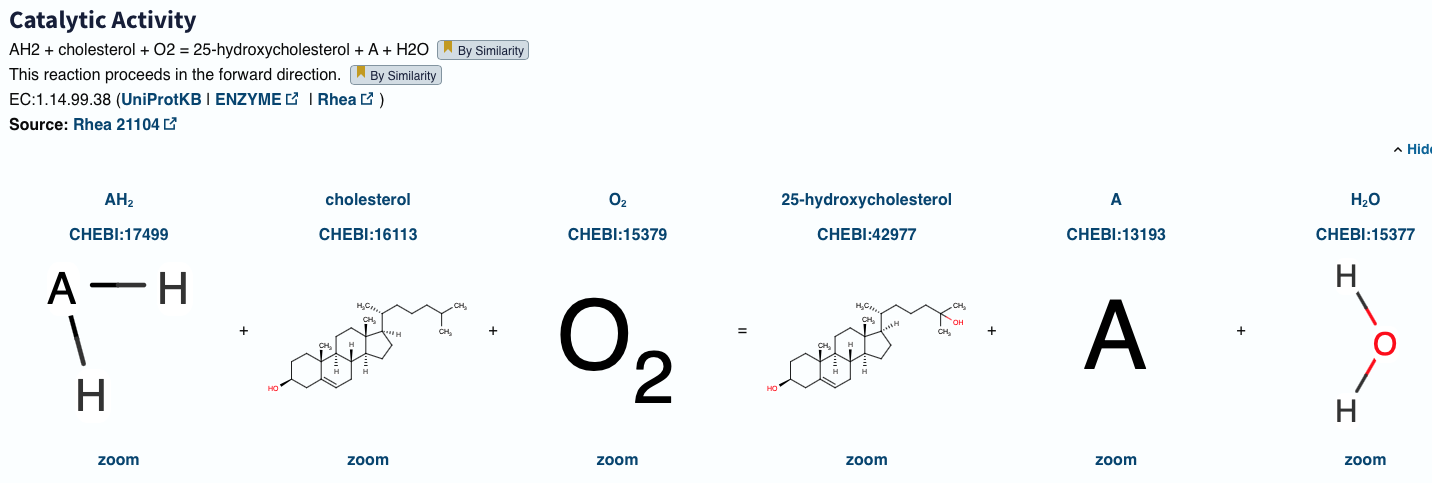

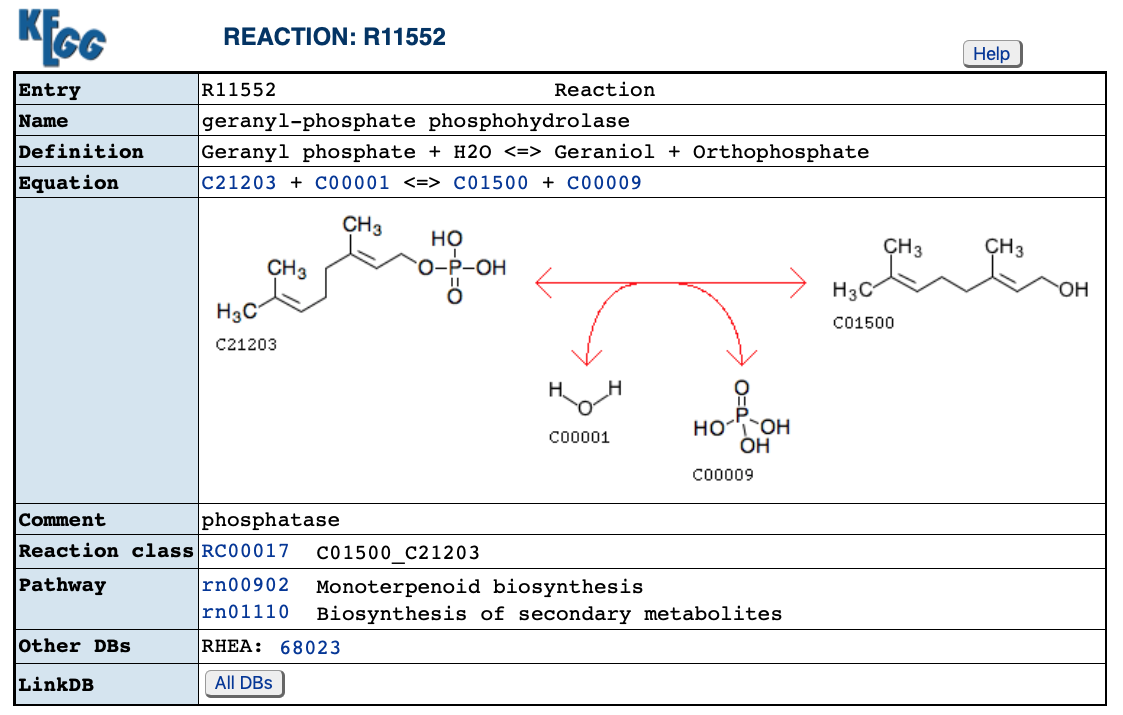

Reaction of the best protein prediction according to BridgIT+:

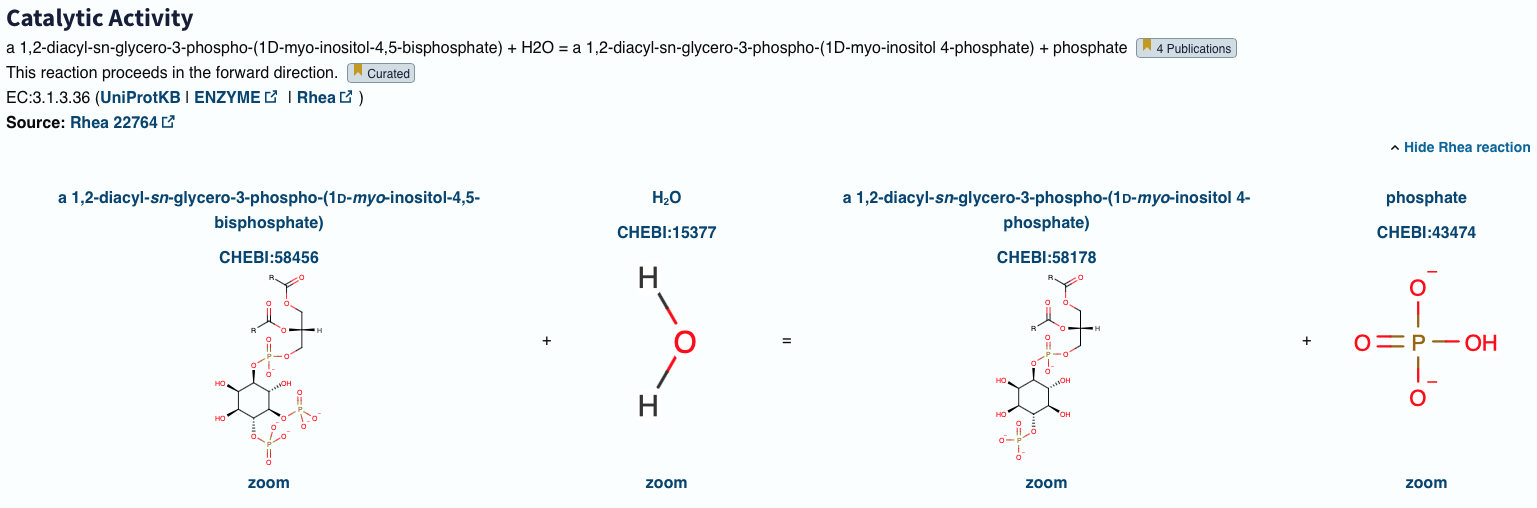

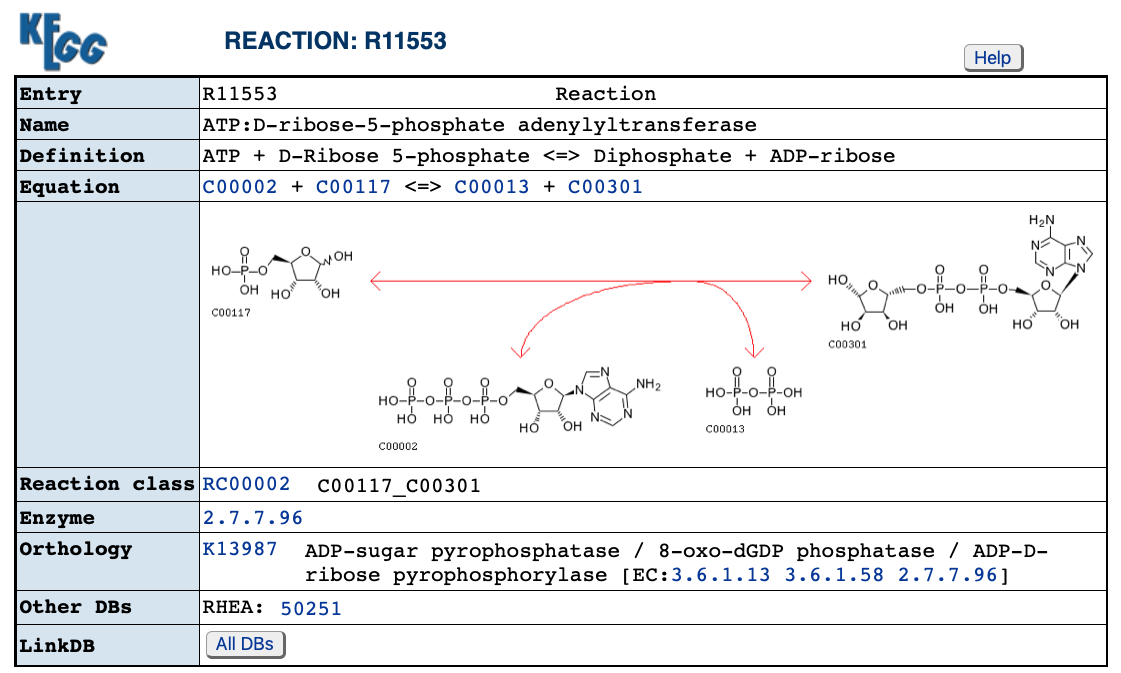

Reaction of the best protein prediction according to BridgIT+:

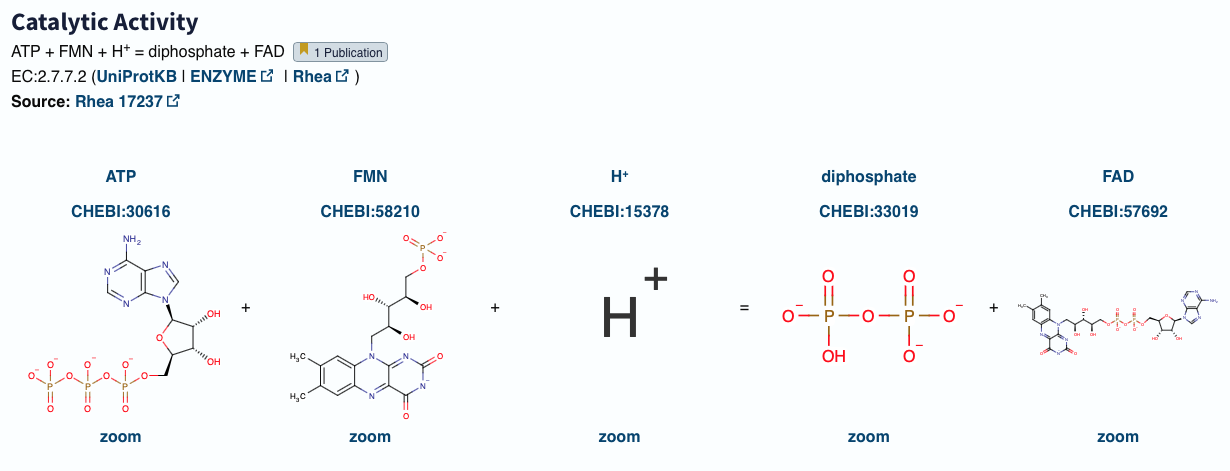

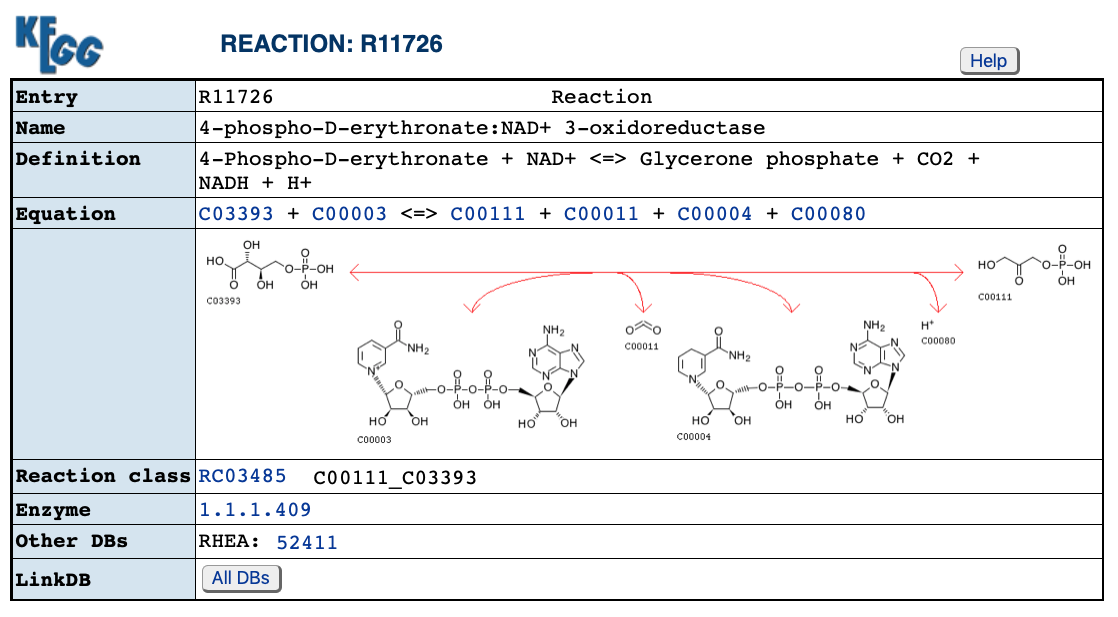

Reaction of the best protein prediction according to BridgIT+:

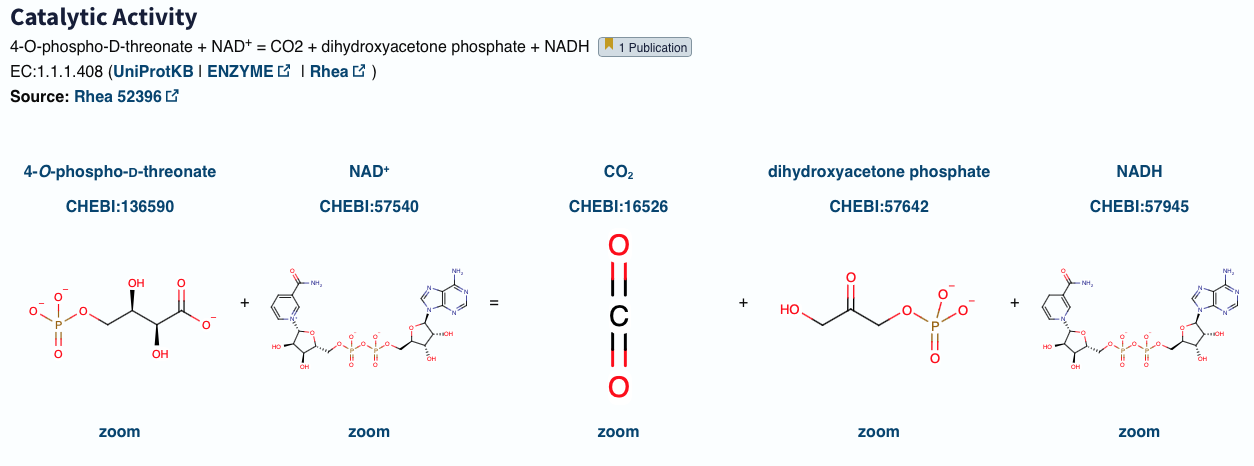

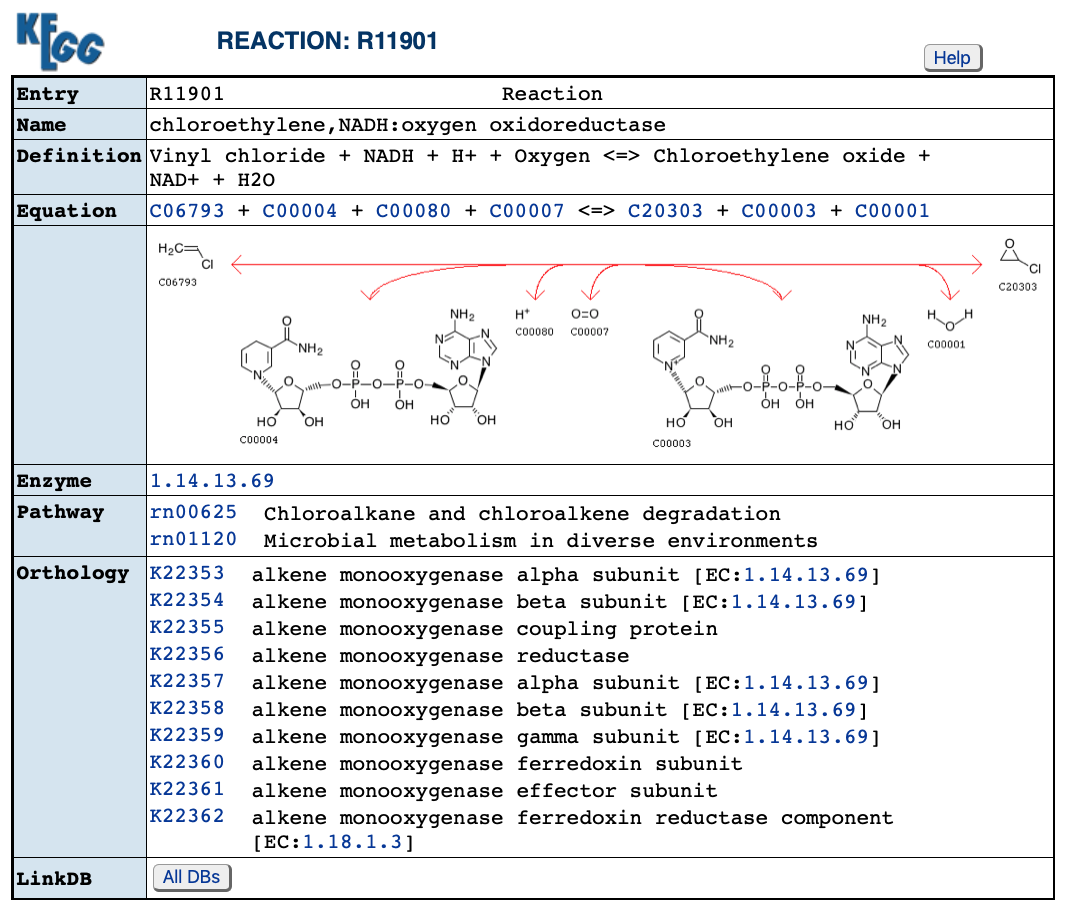

Reaction of the best protein prediction according to BridgIT+:

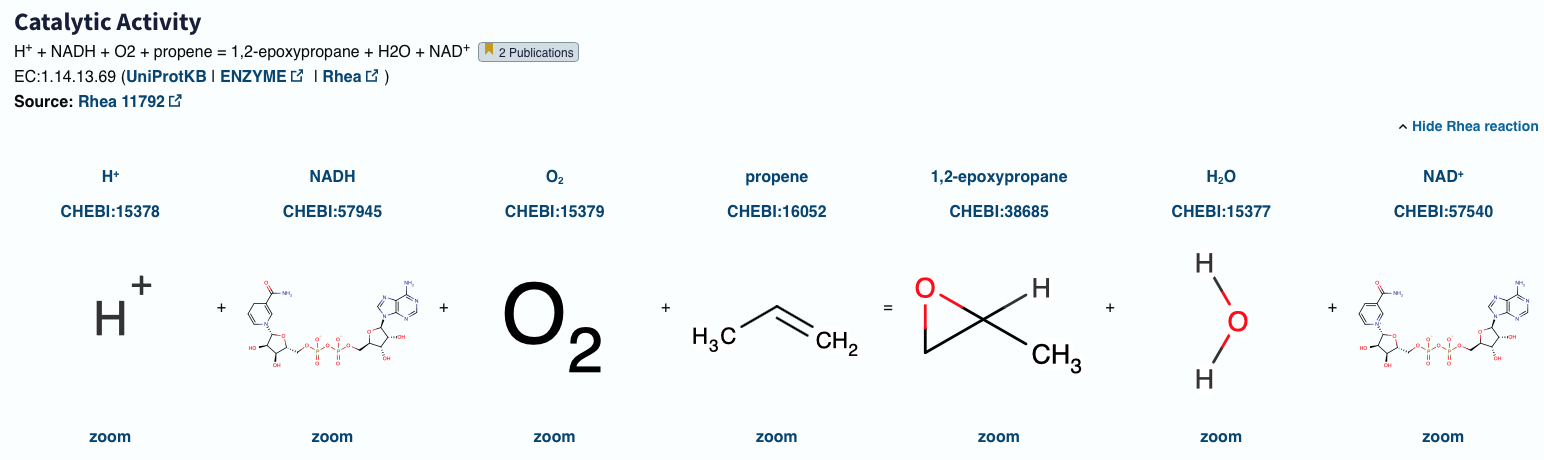

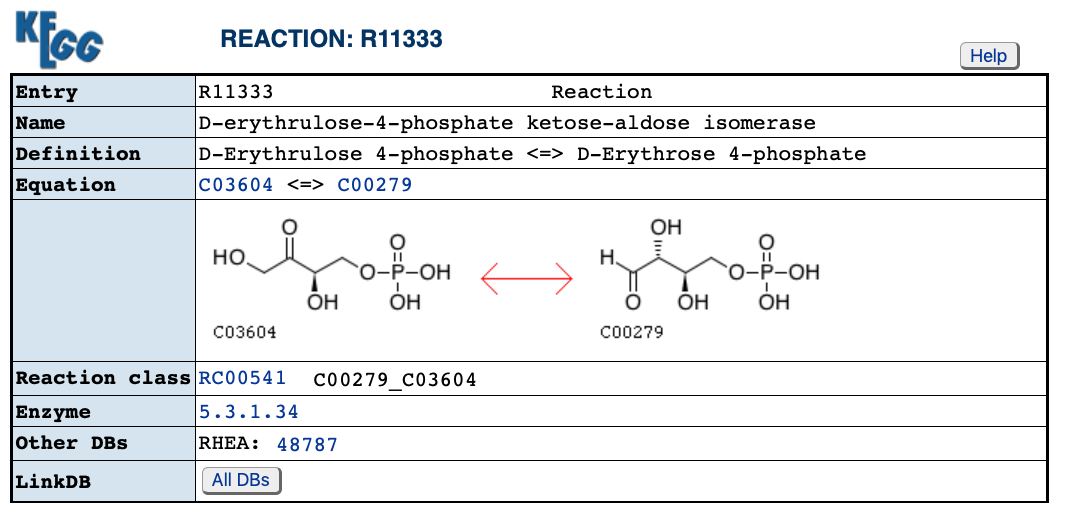

Reaction of the best protein prediction according to BridgIT+:

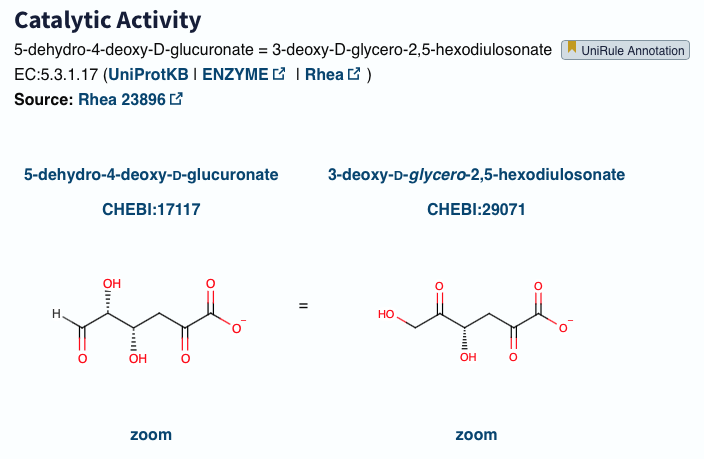

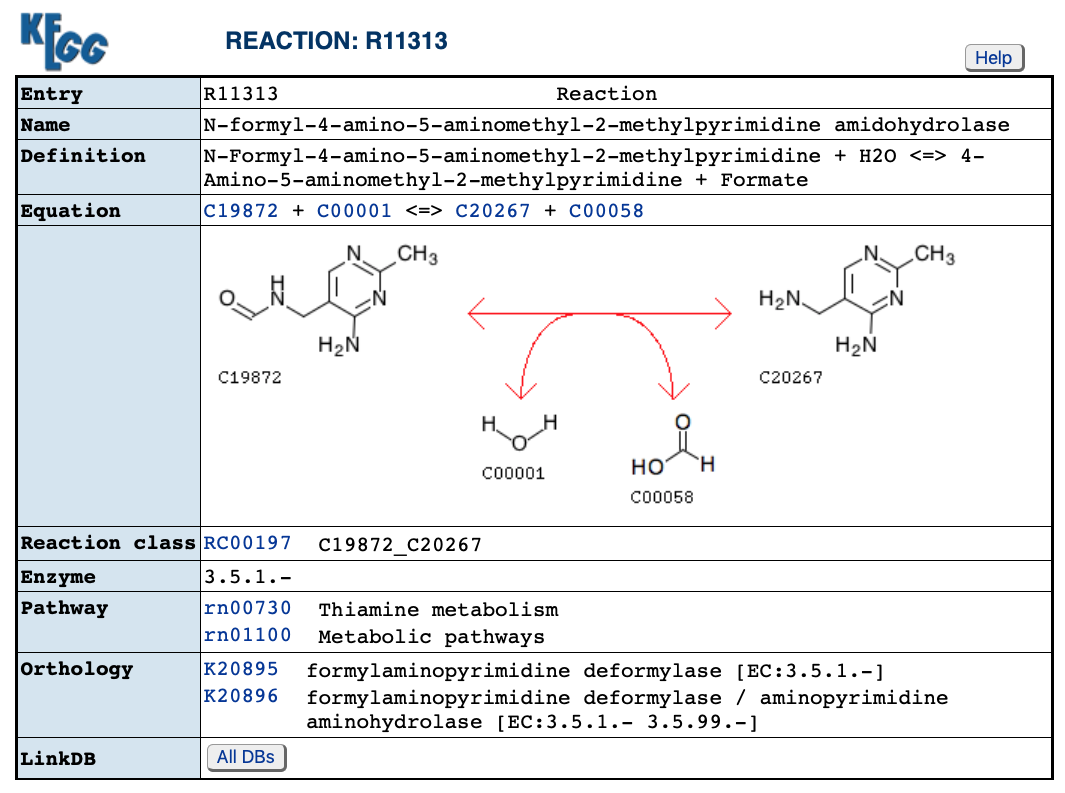

Reaction of the best protein prediction according to BridgIT+:

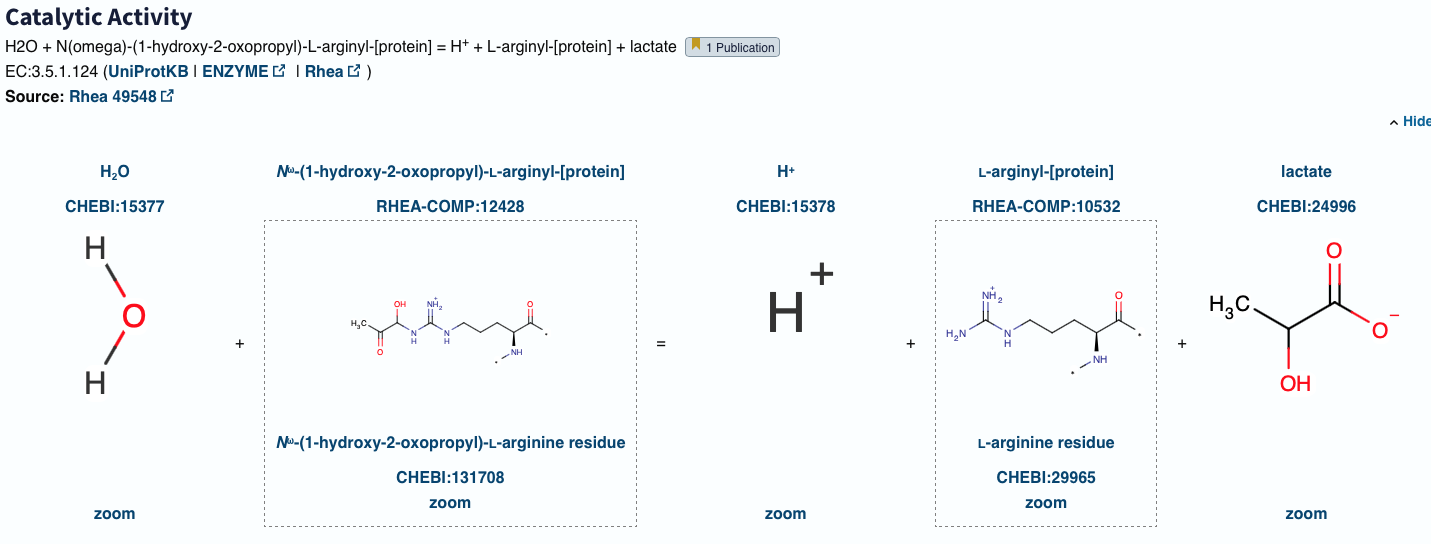

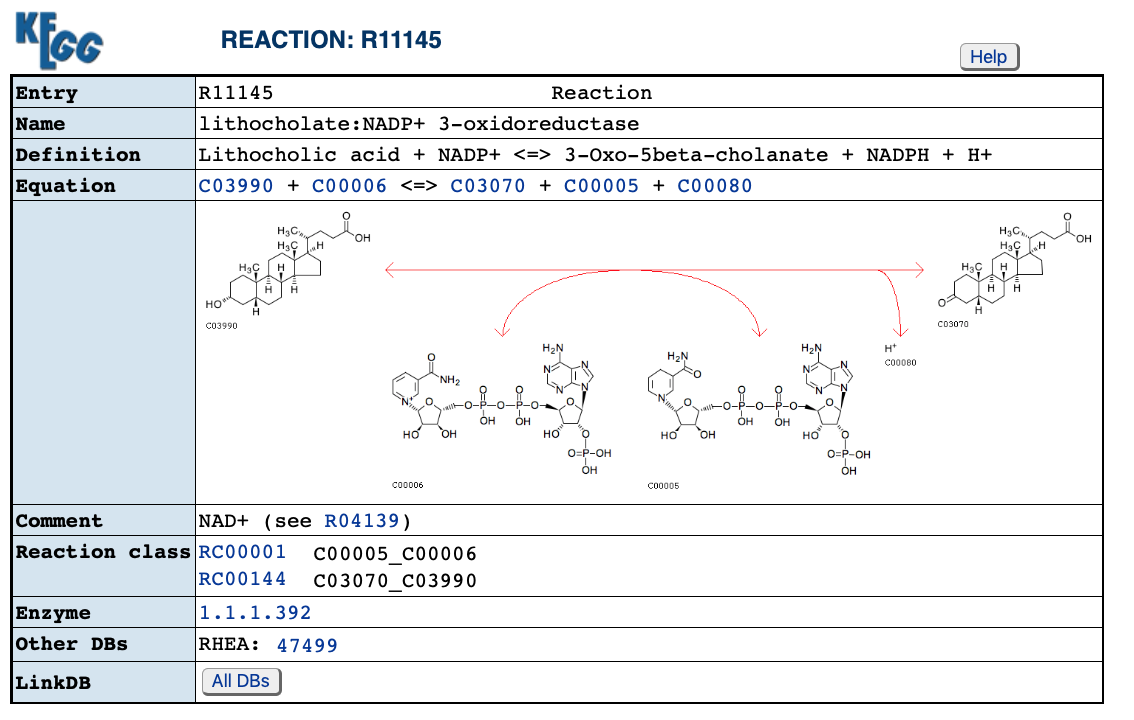

Reaction of the best protein prediction according to BridgIT+:

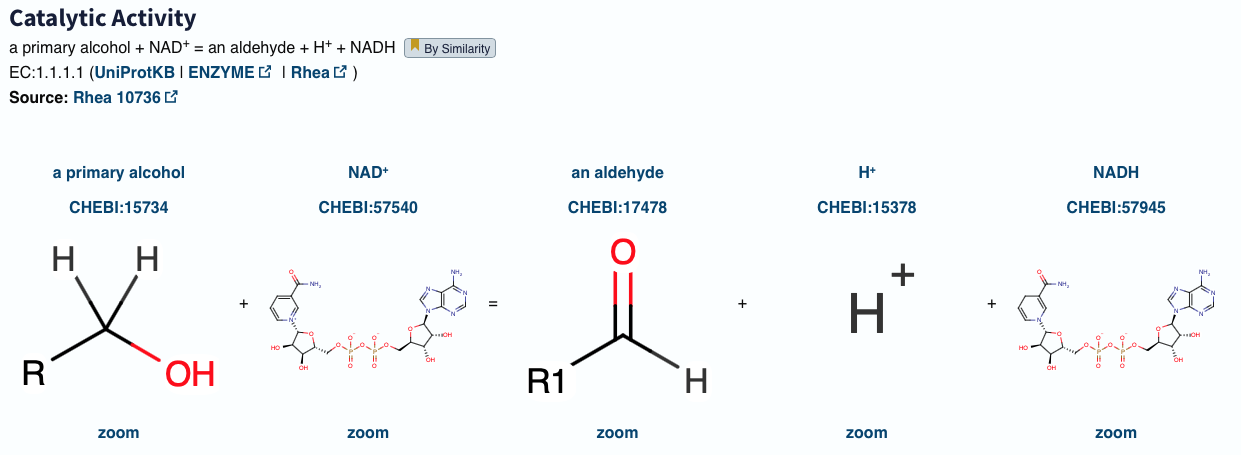

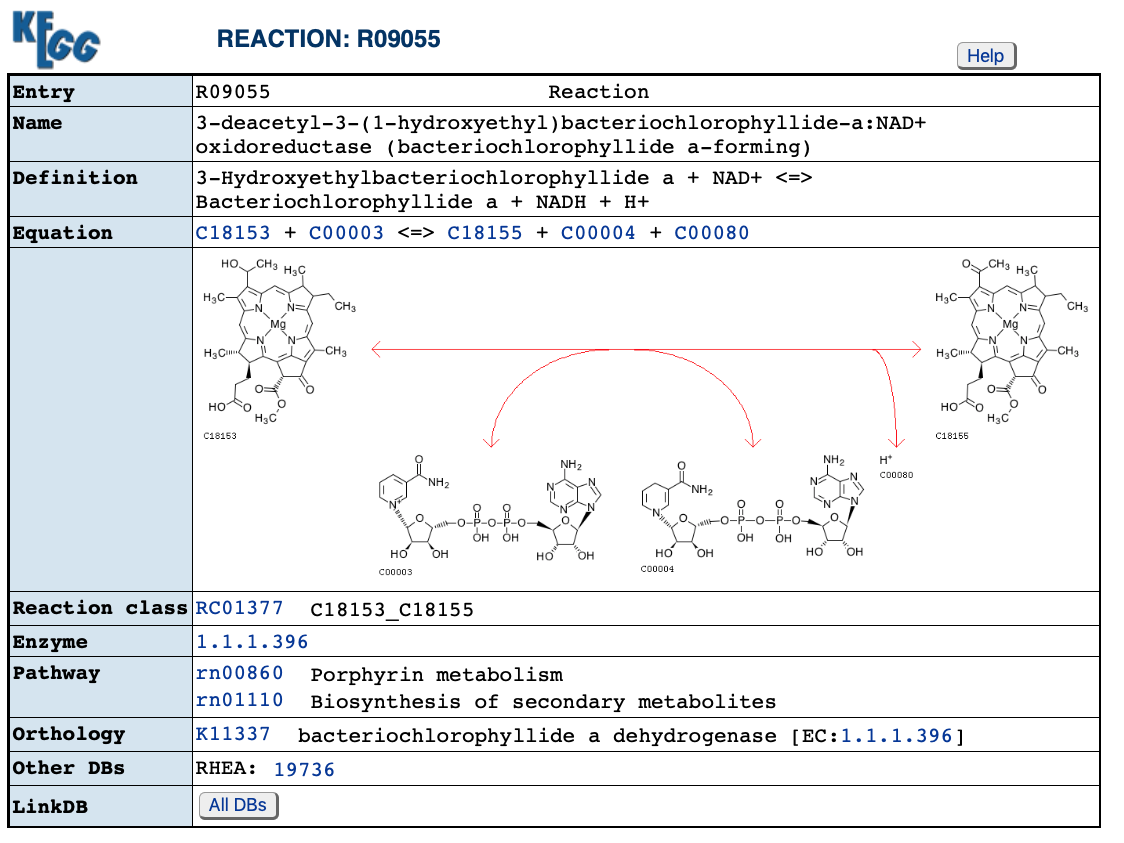

Reaction of the best protein prediction according to BridgIT+:

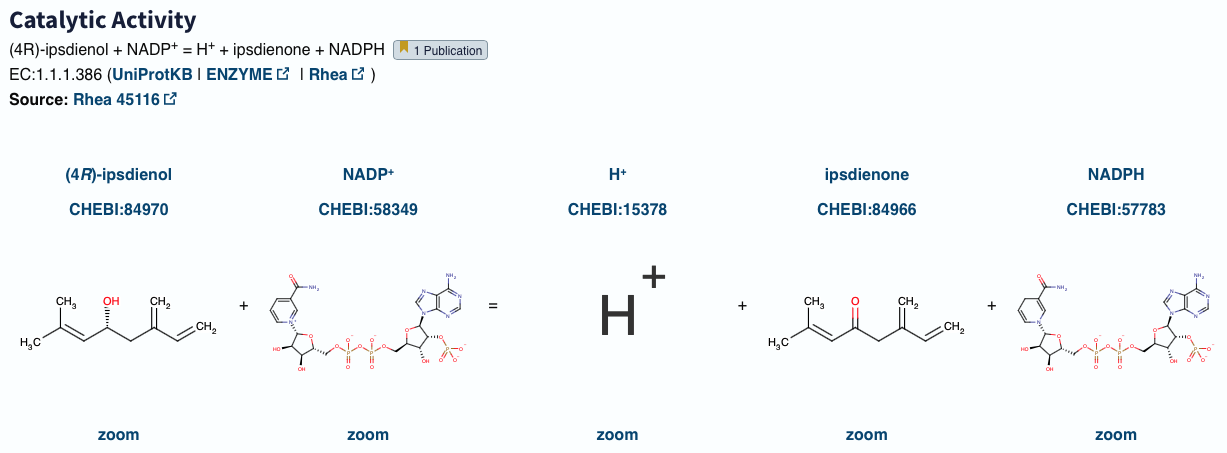
