## Supplementary material 1 for "Enzyme promiscuous profiles for protein sequence and reaction annotation"

**Supplementary 1.**

**Existing protein information resources/ tools:**

**PROSITE:** defines protein domains and sequence patterns of biological significance

**CATH:** Analysis of the structural domains generated by CATH reveals the prominent features of protein secondary structure

**Enzyme 3D:** structural information from Protein Data Bank utilized to facilitate functional annotation beyond only sequence information

**SCOP:** manually provides structural classification to reveal the evolutionary relationships between proteins. Structural information can help to identify binding sites and catalytic residues on proteins further.

**M-CSA:** A specific collection of catalytic residues and detailed processes of catalytic mechanisms can be obtained

**EnzyMine**: A new knowledgebase providing enzymatic reaction features and then link them with sequence and structural annotations

**Protein annotation tools:**

**EnzDP: Uses** functional domain architecture to score the association between domain families and enzyme families (Domain-Enzyme Association Scoring, DEAS). The DEAS score is used to calculate the similarity between proteins, which is then used in clustering procedure, instead of using sequence similarity score.

**Supplementary figures**

A

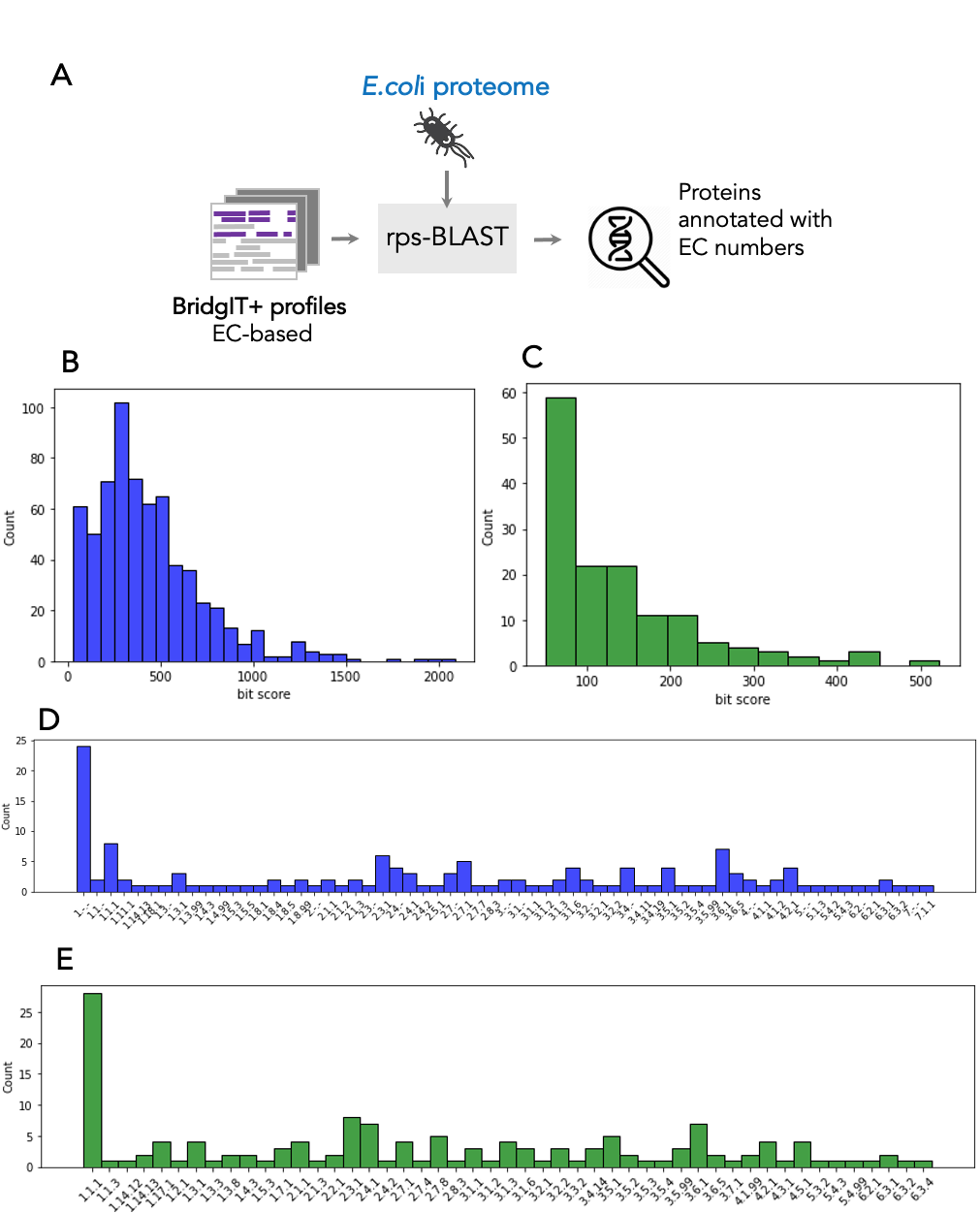

B

Supplementary figure 1. A. EC class distribution for 598 reactions validated with BridgIT+ annotation of E. coli proteome. B. EC class distribution for 144 reactions newly discovered in E. coli proteome with BridgIT+ annotation.
